# Synthesis of arbitrary interference patterns using a single galvanometric mirror, and its application to Structured Illumination Microscopy

**DOI:** 10.1101/2024.02.29.582785

**Authors:** Ke Guo, Abderrahim Boualam, James D Manton, Christopher J Rowlands

## Abstract

We present a new interferometer concept called SWIFT, able to project arbitrary interference patterns constructed from small numbers of plane waves. SWIFT can control each plane wave’s orientation, intensity, polarization and phase using just a single galvanometric mirror. We demonstrate the application of SWIFT to both 2D and 3D Structured Illumination Microscopy, characterizing performance on fluorescent nanoparticles and iFluor 488 phalloidin-stained U-2 OS cells.

---

The projection of optical interference patterns underpins many modern optical measurement techniques, including structured illumination microscopy[1], lattice light-sheet microscopy[2], fringe-projection profilometry[3], spatial frequency-domain imaging[4], parallel STED[5] / RESOLFT[6], and many more techniques besides. In almost all implementations, the interference patterns are created by superimposing two or more beams at different angles; these beams are sensitive to changes in relative path-length, which can lead to pattern instabilities. Common-path interferometers can address this problem; indeed it is possible to think of any image projection system as a complicated common path interferometer. For example, imaging the reflected orders from a diffraction grating [1], spatial light modulator (SLM) [7] or digital multimirror device (DMD) [8, 9] can yield stable interference fringes. Nevertheless this approach is typically power-inefficient due to higher order diffraction losses[8, 10]; additionally, the size and pixelation of SLMs and DMDs limits the field of view and resolution. An alternative approach is to project a small number of point-sources through a lens; by the Fourier transform properties of a lens, these point-sources become plane waves.

In order to control the position, amplitude, polarization and phase of each point-source independently, it is commonly required for every point-source to have a separate actuator for each parameter. Previous solutions using this approach use a suite of galvanometric (“galvo”) mirrors[11], or individually-addressed optical fibers[12]; however, galvo mirrors are costly and complex to synchronize, whereas optical fibers have a low damage threshold, limiting light throughput.

In this work, we present a new method for generating interference patterns in which the location of each source (and thus the orientation of its corresponding plane wave), its polarization, amplitude and phase can all be controlled using a single galvo mirror. Subject to a few constraints, arbitrary interference patterns can be created. The design of this pattern generator (which we call the Synthetic Wide-field Interfering Foci Technique or SWIFT) can be seen in Fig. 1. In brief, a galvo steers a set of mutually-coherent beams to illuminate a column of mirrors. Each mirror steers its beam to an arbitrary lens in a lens array (e.g. the two mirror-lens pairs highlighted in red and green in Fig. 1), thus forming a series of foci at the back aperture of a large-aperture lens. The rays from each focus are collimated by this large-aperture lens, forming a desired pattern as they interfere. Because the angle of each mirror ultimately controls which lens array element is illuminated (and therefore the angle of the interfering beam), changing the mirror tilts changes the resulting interference pattern. To change the interference pattern dynamically, another column of mirrors is placed adjacent to the first, each of which steers the incident light to a different lens; the galvo is used to illuminate the new column of mirrors, ultimately creating a different interference pattern. More subtly, because the galvo is positioned conjugate to the back aperture of the lens, by stepping a small distance it can apply a small phase shift to each beam, proportional to its horizontal tilt; thus, the phase of the interference pattern can controlled. Furthermore, by placing a quarter wave plate and an attenuator over each mirror, the amplitude and polarization of every point source can be individually controlled. Compared to direct projection of a grating, SLM or DMD, the system is achromatic, does not suffer from higher order diffraction and does not have limits on field-of-view and/or resolution caused by finite numbers of pixels. To demonstrate the utility of SWIFT, we use it to generate the illumination patterns for a high-speed Structured Illumination Microscope (SIM) [1]. SIM is an optical super-resolution microscopy technique that uses the Moiré effect to increase the resolution up to twice the classical diffraction limit. It works by projecting a series of very finely-spaced fringes onto the sample, traditionally at three different orientations and with three [1] or five [13] different phases per fringe pattern. We show that SWIFT can be used to generate both the 2-beam and 3-beam interference patterns required for 2D and 3D SIM, and switch between all the orientation and phase combinations by controlling only the galvo. The quality of the generated patterns are highlighted by successful 2D SIM imaging of fluorescent nanoparticles at 89 fps (980 fps raw frame rate), and 3D imaging of iFluor 488 phalloidin-stained U-2 OS cells.

**Fig. 1.**
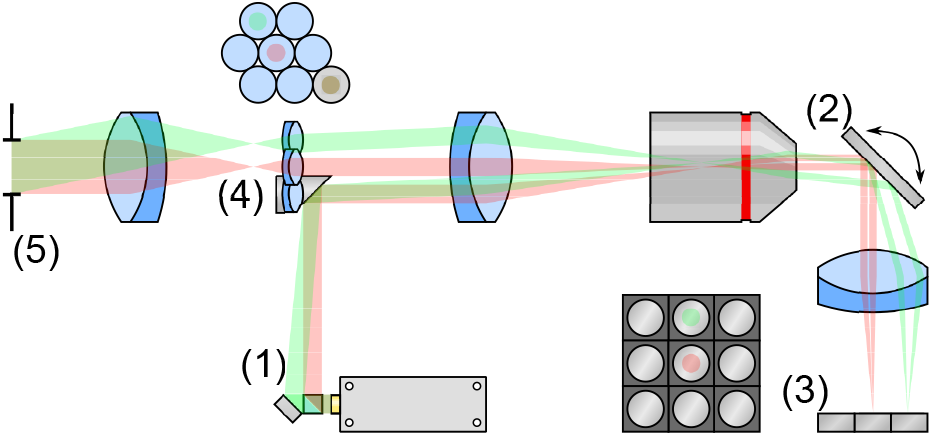
SWIFT design. Light from a laser at (1) is split into multiple beams (only two are illustrated for clarity in red and green), which are reflected onto the main optical path via a mirror. The beams are imaged and demagnified onto a galvo mirror at (2). The galvo steers each beam through an achromatic lens onto an array of miniature mirrors mounted on small kinematic platforms at (3). Each of the platforms can be controlled independently, to reflect the beam to an arbitrary location on the galvo. The galvo is in a conjugate plane to a miniaturized lens array at (4); the purpose of the miniature mirrors is therefore to steer the retroreflected beams to any of the miniature lenses in the array. The focal plane of this lens array lies at the back focal plane of a achromatic lens, which forms the desired interference pattern at (5). Controlling the galvo at (2) allows the user to illuminate a different column of miniature mirrors, which in turn allow a different set of lenses to be illuminated. In this way, the user can pick between a collection of pre-set arbitrary interference patterns at will.

## 1 Results

### 1.1 Design

SWIFT consists of five main parts: a <u>beamsplitter unit</u>, a <u>4f demagnifier</u>, a <u>galvo mirror assembly</u>, a <u>miniature mirror array</u>, and finally a <u>miniature lens array</u>. Beamsplitter units can be constructed from a diffractive element, as is typically done for SIM, but to achieve achromaticity and minimise loss of light. We designed a beamsplitter unit using a series of achromatic half-wave plates and polarizing beamsplitters (see Fig. 2 and Supplementary Fig. 2). The power of each beam is controlled by the halfwave plates, which change the beam’s polarization and thus the fraction reflected from the beamsplitters. A subsequent set of pick-off mirrors allow the beams to be steered independently while passing close to each other. The unit was assembled on a custom-machined baseplate using small FiberBench components for stability and cost-efficiency.

**Fig. 2.**
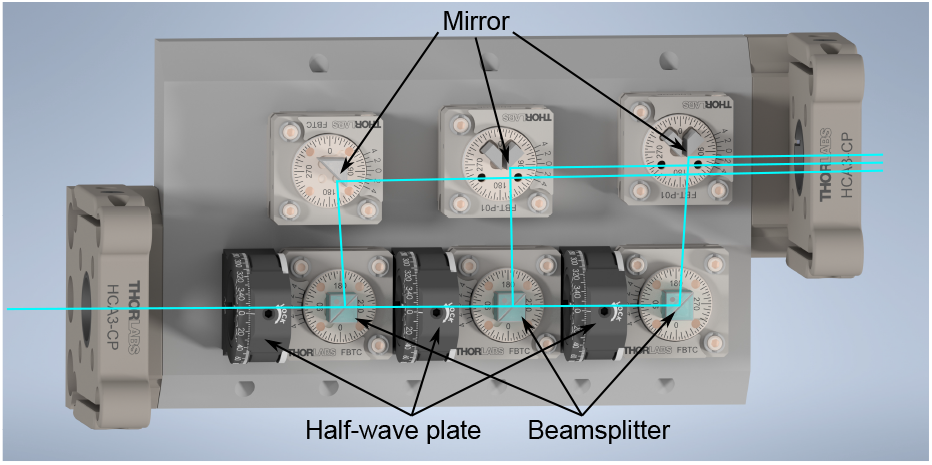
Beamsplitter array. The cyan beam entering on the bottom left passes through three sets of half-wave plates and polarizers, which control the relative power of the reflected beams. These beams (each nominally s-polarized) pass through a Glan-Laser prism (not visible) to improve the polarization purity.

After the beamsplitter unit, the beams strike a pick-off mirror and are imaged onto the galvo using a telecentric 4f optical system with a substantial demagnification (4 *×* in our case). The galvo reflects the beams to their respective mirrors, addressing a column of mini mirrors at a time. Each mini mirror is mounted on its own kinematic platform with tilt adjustments and a screw thread for axial displacement in order to match path-lengths (see Supplementary Fig. 4). The mini mirrors reflect the light back towards the galvo and then to the miniature lens array. Because the galvo surface is conjugate to the lens array, the tilt of each mini mirror (controlled by its kinematic mount) determines which lens the beam passes through. Attenuation and polarization control can be performed on a per-mirror basis by placing a neutral density filter and a polymer quarter-wave plate in front; the light passes the wave plate twice, and thus it acts as a half-wave plate, rotation of which controls the linear polarization axis orientation.

The focus of the lens array is at the back focal plane of a large lens; therefore, each focus is converted to an angled plane wave, all of which interfere at the front focal plane to produce the desired interference pattern. Since the coarse position of the galvo determines the column of the mini mirrors addressed by the beams, switching between columns allows different interference patterns to be created, with arbitrary combinations of angle, intensity and polarization, albeit with the requirement that these parameters must be determined ahead of time. Furthermore, when moving the galvo a smaller distance such that beams still address the same column of mirrors, the path length of one beam can be changed relative to another. Consequently, a phase shift is applied to the interference pattern, without changing the structure of the pattern itself. The ratio of the phase shift to the change in the galvo angle is proportional to the horizontal distance between the relevant lenses.

The galvo stability requirements are dictated by the size of the lens array and the aforementioned demagnification ratio; the galvo is conjugate to the lens array and therefore to maintain a desired phase relationship between the outermost beams, the galvo must not jitter excessively. For example, to maintain a *λ/*10 stability at 488nm, with a 2 mm separation between the beams on the galvo, the galvo should be stable to arctan(48.8 nm*/*2 mm) ≈24 μrad. Fortunately, the Dynaxis 3s galvo selected for the prototype boasts better than 1 μrad repeatability.

### 1.2 Performance

To assess interference pattern generated by SWIFT, we imaged the centre of the interference pattern using a separate home-built 4*×* microscope. Assessments were performed on three types of interference patterns: a two-beam interference pattern, a hexagonal three-beam pattern and a linear three-beam pattern. Figure 3(a-b) show a measured two-beam interference pattern generated by overlapping the top and bottom beams originating from the positions on the back aperture highlighted in the inset. By fitting the image to a 2D sinusoidal function with an offset, we obtain the visibility (*V* = (*I*_max_ *−I*_min_)*/*(*I*_max_ + *I*_min_)) of fringes to be 0.92. Similar results can be obtained from the other two orientations. A hexagonal pattern was generated by overlapping three beams through the outer lenses, as shown in Figure 3(c-d). As a horizontal offset between the lenses is necessary for the phase shift, the lens array in this case was rotated by 90^*°*^ from the two beam interference setting. Finally, Figure 3(e-f) show the three-beam interference pattern commonly used for 3D SIM [13]. Because this pattern is periodic in z as well, we also measured the intensity distribution as a function of axial position in Figure 3(g-h). To do so, the objective of the home-built microscope was moved along the optical axis using a high-precision motorized translation stage. The measured images show high contrast fringes similar to the expected three beam interference pattern, except for a shift in the vertical direction in Figure 3(g) which is attributed to misalignment between the SWIFT optical axis and the direction (z axis) along which the camera was moved.

**Fig. 3.**
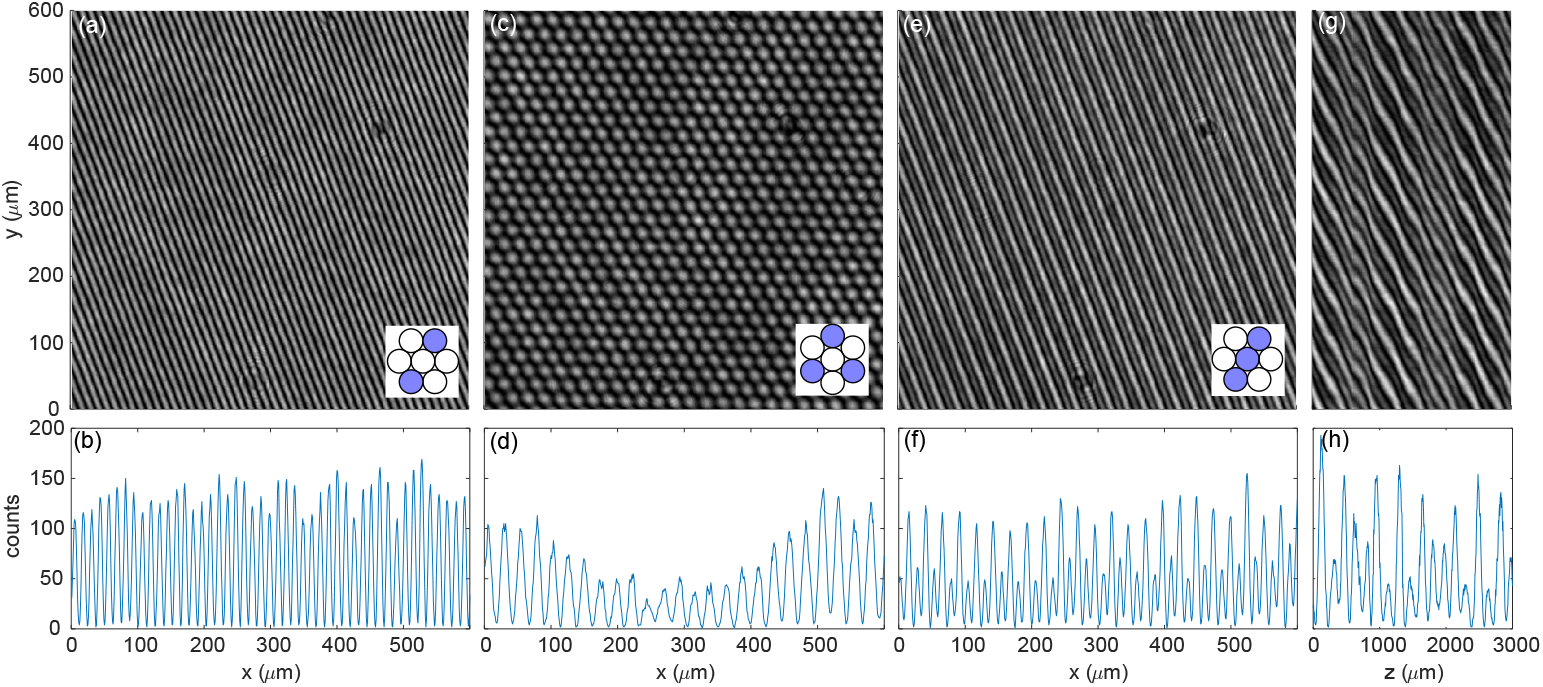
Centre of the interference patterns near an intermediate image plane measured using a 4 *×* microscope. (a) Two-beam interference pattern, (c-d) Hexagonal interference pattern. (e-h) Three-beam interference pattern. (g) is obtained by combining y-cuts in the middle of 601 camera images of the three-beam interference pattern at different axial positions. (b,d,f,h) are horizontal cuts of (a,c,e,g) at *y* = 0. The insets illustrate the orientation of the lens array, with the lenses used highlighted in blue.

**Fig. 4.**
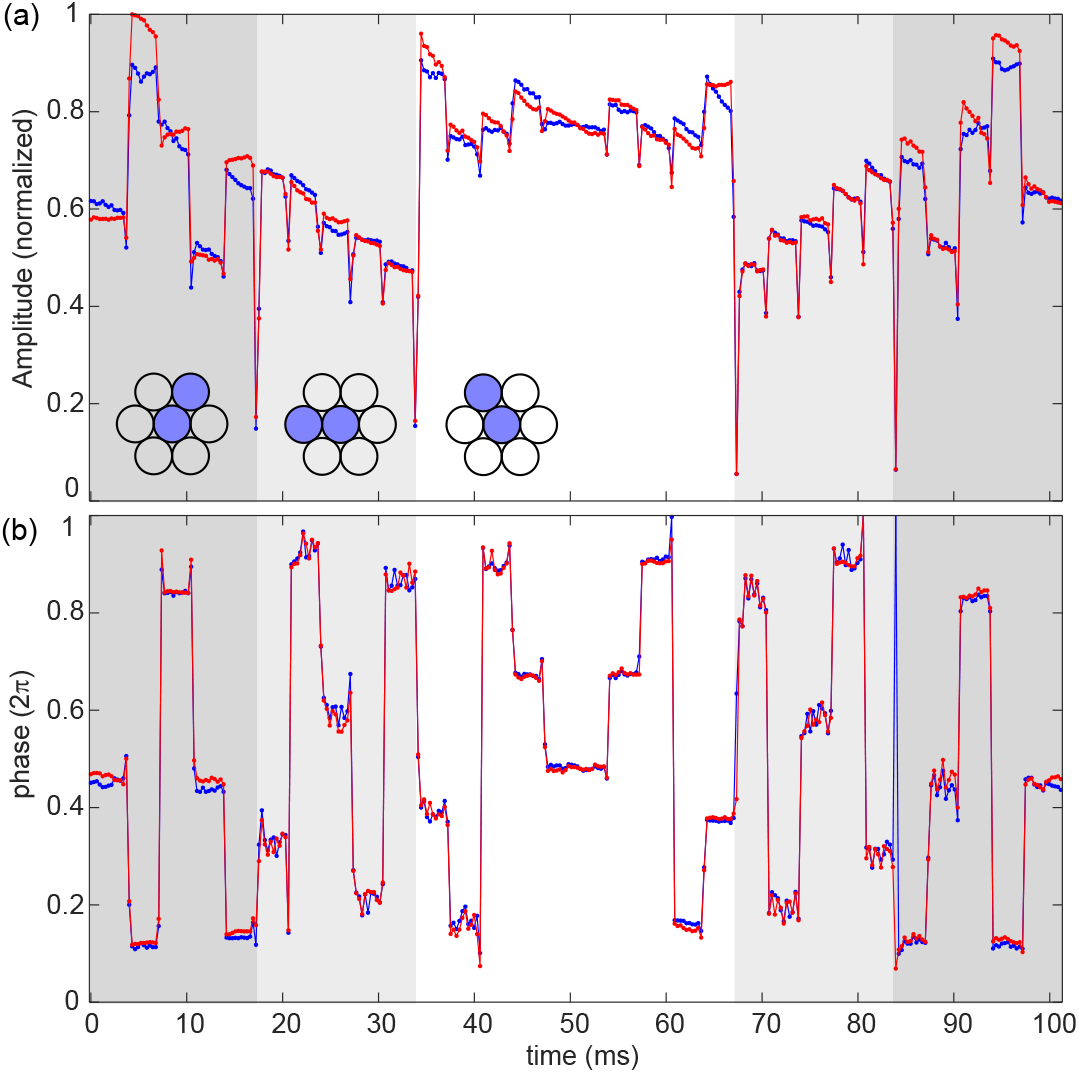
(a) Maximum amplitude of the correlation and (b) the corresponding phase obtained from two repeated sequences (red and blue) of fluorescent images taken at 3252 fps while switching the galvo with the same setting at 300 Hz. Background colours indicate the three different orientations of the fringes and insets illustrate the corresponding mini lenses. The phase of the two image sequences match with fluctuations of a few percent of 2*π*. Laser power: 100 mW. Exposure time: 300 μs.

We assessed the speed at which the interference pattern could be changed by recording high-frame-rate fluorescence images at 3252 fps. The interference patterns were projected into a 100*×* microscope which was used for SIM measurements in the following section. A layer of 200 nm fluorescent beads was used to reveal the interference pattern. To reach the high frame rate, the camera’s Region of Interest (ROI) was set to 65 μm *×* 31 μm in the centre of the sensor. In order to resolve the pattern with the microscope, we use the interference between the top and middle beams using the lenses highlighted in Fig. 4. The galvo was turned back and forth at 300 Hz to switch between the three orientations, with five phase steps per orientation. The phase steps were set to be slightly larger than the steps used for 2-beam and 3-beam SIM.

To assess the quality of the interference patterns, we correlated the Fourier transform of each image with the average of all images. The amplitude, phase and position of the maximum of the correlation reflect the visibility, phase and orientation of the interference patterns respectively. When the mirror reached a stable state, the amplitude of the correlation stayed at a plateau. As the mirror switched, the amplitude of the correlation dropped significantly during the transition frame(s), with concomitant uncertainty in the phase value. Fig. 4 shows the maximum amplitude of the correlation and the corresponding phase obtained from two sequences of images with the same mirror settings. The phase transitions commonly happened within one frame, while the transition between different angles took up to 3 frames, i.e. 0.9 ms. As the mirror rotation for phase shift was less than 1 % of the rotation for the orientation transition, the results indicate that the actual phase transition time can be less than 10 μs. The experiment was performed twice, with the phase of the two image sequences (shown in blue and red) matching well. This indicates that the phase is repeatable and stable, with fluctuations of just a few percent of the full 2*π* range, albeit slightly higher than the theoretical stability of the galvo. We attribute this small decrease in performance to electronic noise in the the galvo driver.

### 1.3 Structured Illumination Microscopy (SIM)

We chose to demonstrate the utility of SWIFT by using the generated interference fringes as illumination patterns for 2D and 3D SIM. SIM is a challenging application, demanding good phase stability, high speed, good optical throughput, and high interference contrast. For 2D SIM, we illuminated with just two beams; the beamsplitter unit was adjusted to reduce the power of the middle beam to near-zero, and a custom-made beam blocker placed in front of the middle row of the mirror array was used to eliminate any residual intensity.

The imaging speed for 2D SIM is commonly limited by the switching time between different imaging patterns, the minimum exposure time to obtain enough fluorescent signal, and the speed of the camera. SWIFT is capable of high speed SIM imaging, with transition speed up to approximately 50000 frames per second for phase shift and 500 frames per second for angle change, for a 65 μm *×* 65 μm ROI, assuming perfect synchronization between the mirror and the camera. To demonstrate this capability, we imaged a layer of 100 nm fluorescent nanoparticles deposited under a layer of UV-cured adhesive which was used to match the refractive index of the immersion oil. As the camera frame rate is limited by the number of pixel rows, we used only the centre 1000*×*200 pixels to achieve a frame rate of 980 fps. The galvo signal was synchronized with the camera trigger to repeatedly move the mirror to one of 9 positions, specifically 3 phase steps for 3 orientations. As the direction of motion alternated, the mirror position was unchanged during the first frame of each imaging cycle. To accommodate the longer switching time (approximately 1 ms) between different orientations, the mirror was programmed to remain for two frames in the first phase of the second and third orientations. Consequently, each cycle took 11 raw camera frames, yielding a final imaging rate of 89 fps. With the first frames after the orientation change discarded, 9 out of the 11 frames were processed using a home-made MATLAB program based on open source algorithms [14, 15] to produce a super-resolution image. A theoretical PSF assuming NA=1.48 was used for reconstruction. When operating at a high frame rate, we observed microscale motion of the sample relative to both the interference pattern and the camera which can cause strong artefacts in the reconstructed image. Thus, the raw camera images were pre-processed using a sub-pixel image registration algorithm [16] before reconstruction. Figure. 5 shows an example of a reconstructed image along with the corresponding wide-field image with the theoretical PSF deconvolved. The reconstructed SIM image shows a clear improvement in resolution which is sufficient to resolve beads with a separation of 100 nm, such as the two marked by arrows in Figure. 5(d). Figure. 5(d-e) shows a comparison of reconstructed SIM images with (d) and without (e) compensating for camera motion. The stronger artefacts in (e) demonstrate the degradation of reconstructed image quality caused by the camera motion, but also that this degradation can be resolved by pre-processing.

**Fig. 5.**
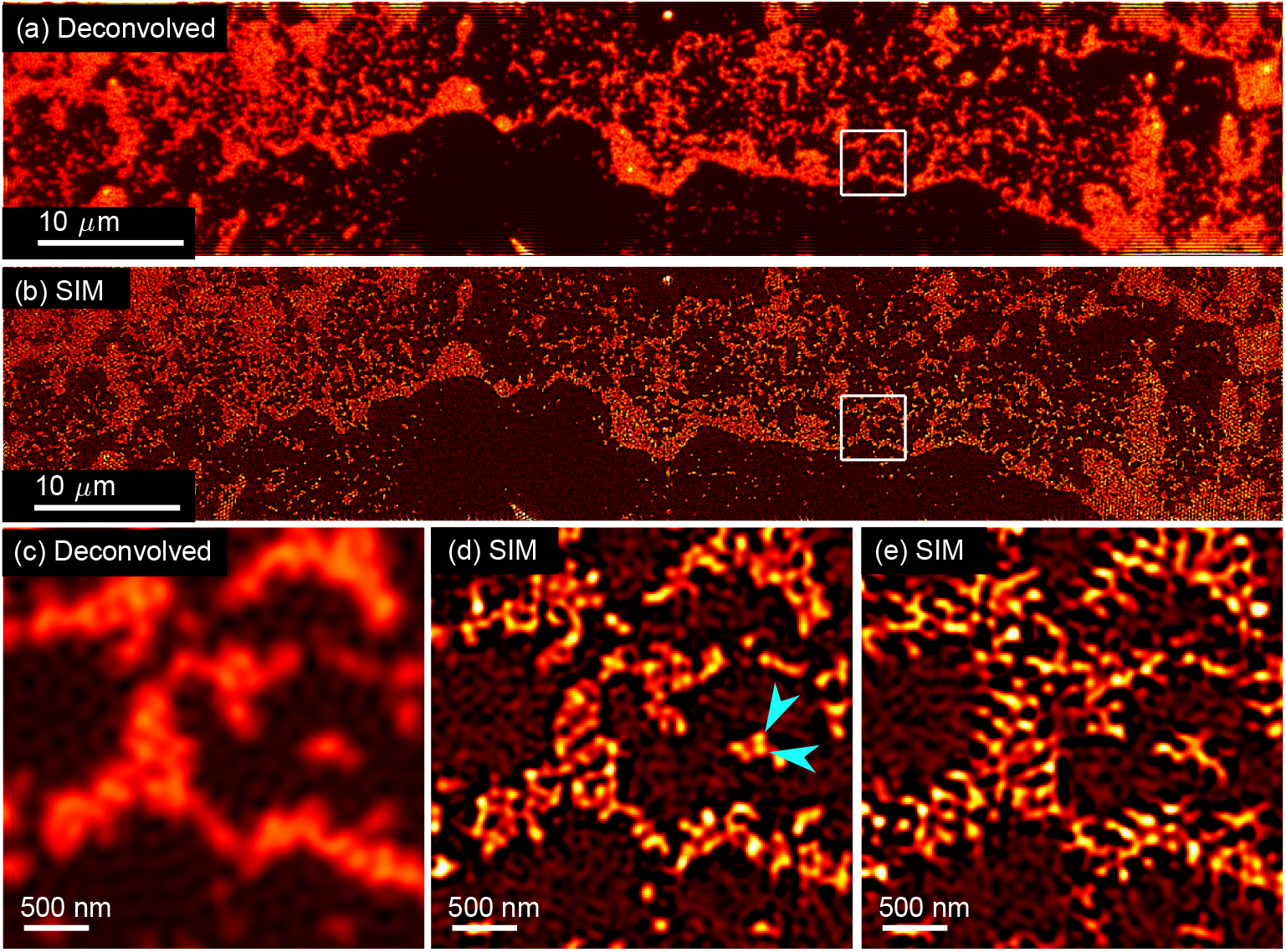
Images of 100 nm fluorescent beads. (a) A widefield image obtained by averaging 9 SIM frames and deconvolving the theoretical PSF. Camera motion was removed in pre-processing. (b) Reconstructed SIM image with camera motion removed. (c-d) Zoomed sections of (a-b) at the same position. The arrows in (d) point to two beads with a vertical peak separation of 100 nm. (e) Reconstructed SIM image of the same regime as (c-d) without removing the camera motion, showing strong artefacts. Exposure time: 1 ms per frame, final super-resolved frame rate: 89 fps. Laser power: 400 mW.

We applied 3D SIM imaging to iFluor 488 phalloidin-stained U-2 OS cells with 3-beam illumination. The beamsplitter unit was adjusted to optimize interference contrast. To obtain steps the z direction, we moved the objective using a stepper motor focus drive with a step size of approximately 130 nm. At each position, 15 frames were taken with 3 orientations and 5 phase steps. Figure 6 shows a comparison of a deconvolved widefield image and a SIM image. The SIM image shows clearly improved resolution in both lateral and axial directions.

**Fig. 6.**
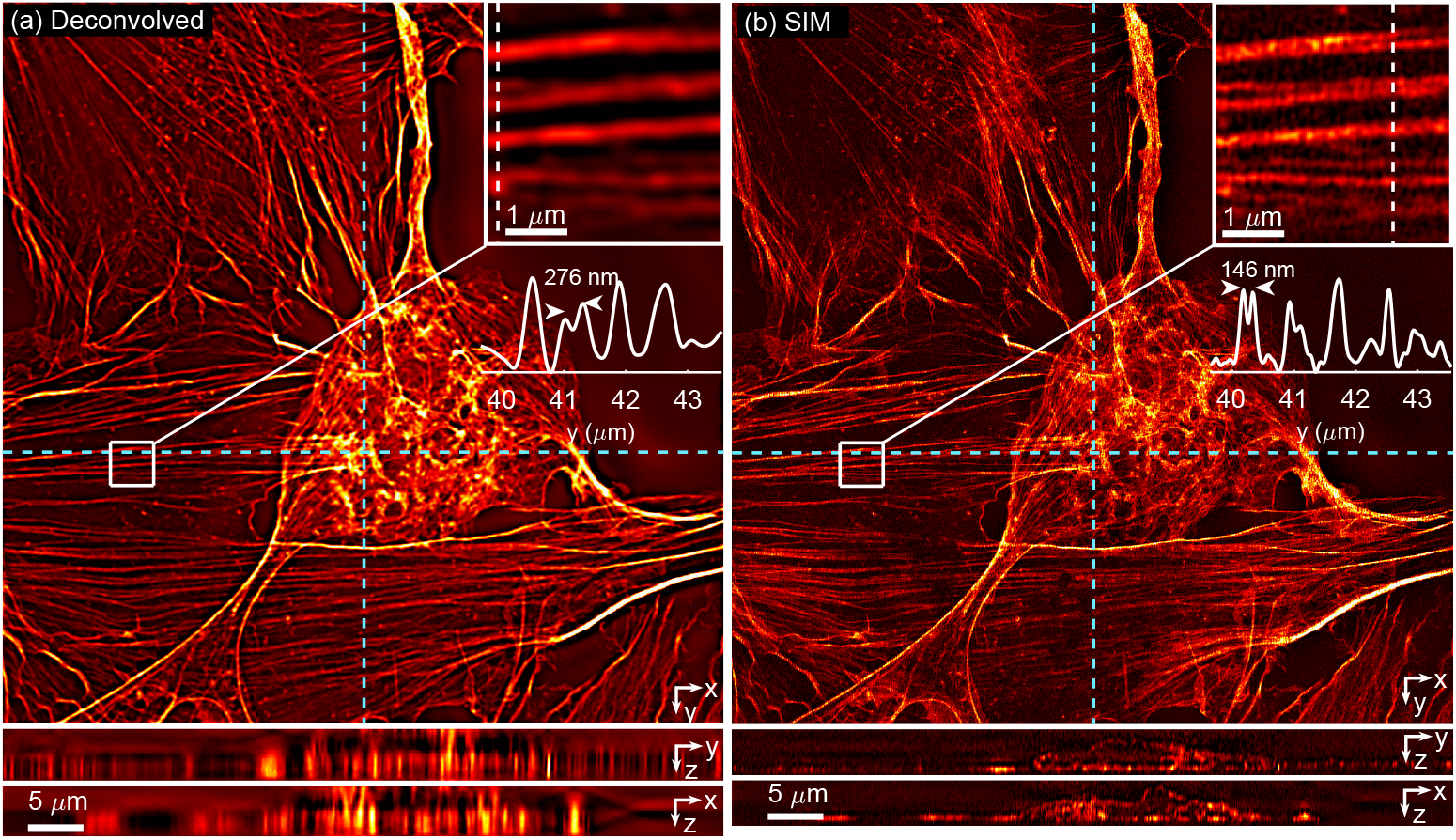
3D SIM imaging of U-2 OS cells stained with iFluor 488 phalloidin. (a) Sections of a 3D widefield image after deconvolving the theoretical PSF. YZ and XZ projections below correspond to the locations marked with the cyan dotted lines. An inset image shows the diffraction-limited imaging performance in a selected region with a cut along the white dotted line. (b) Sections of a reconstructed 3D SIM image. YZ, XZ projections and inset image are taken from the same locations as (a), with resolution improved in both the lateral and axial directions. Exposure time: 199 ms per frame. Laser power: 20 mW.

### 1.4 Discussion

The SWIFT concept contains a number of subtleties that warrant discussion. The first is the 4*×* demagnification between the lens array and the galvo, which was chosen as a trade-off between the system stability and practicality. Since the galvo and lens array are conjugate to each other, a stronger demagnification will make for a smaller image on the galvo and thus increased phase stability against the jitter of the galvo. Nevertheless, this comes at the cost of larger spots on the miniature mirror array, potentially requiring the mirror array to be impractically large.

Next is the choice of parameters for the lens array, the purpose of which is to maintain a small spot size at the back aperture of the final lens, thereby increasing the diameter of the beams at the plane of the interference pattern and thus increasing the patterned area to practical levels. There are no theoretical restrictions on the location, focal length or diameter of the miniature lenses, since a decrease in diameter can be compensated by a proportional reduction in focal length, but on a practical level this reduction in lens diameter cannot be applied excessively when considering the stability of the kinematic mounts in the mirror array. These mounts determine the positions where the beams strike the lens array; smaller lenses would require more accurate kinematic mount alignment and greater stability, as well as requiring that the beam size must be smaller than the lens diameter. It should be noted that the lenses need not be in any regular pattern; arbitrary patterns can be created by positioning individual lenses, by 3D-printing[17], by single-point diamond turning or photolithography[18].

The final subtlety is the nature of the interference pattern phase shift by small motions of the galvo. To explain this effect one must observe that, as the galvo rotates, the focus on the mirror array shifts slightly. If the mirror has a horizontal tilt component, the focus shift will incur a small change in path length, proportional to the angle of the tilt. The overall result is to shift the whole pattern left or right. This also highlights a key fact, that if the lenses illuminated in the array are located have the same horizontal (x) coordinate, no relative phase shift is possible. This could be overcome if the galvo were capable of tilting along both X and Y, but a more practical approach for many applications is to just rotate the lens array slightly.

Aside from design considerations, there are limitations too. For example, while SWIFT can hypothetically include infinite combinations of interference sources, in practice the number of sources is limited to the number of laser beams generated in the beam-splitter unit or the number of mirrors in each column; in addition, the source locations are predefined by the lens array (although as noted above, the number of lenses could theoretically be increased substantially over this prototype). The intensity and polarization of each beam is controlled by the corresponding attenuator and waveplate over each mirror in the array, so in reality, only the phase of the interference pattern may be changed on-the-fly. For many applications though, this an acceptable limitation.

Despite the above constraints, SWIFT can generate useful interference patterns for many applications. We demonstrate its use in 2D and 3D SIM. A major advantage of SWIFT is its high speed, which is important for capturing super-resolved dynamic biological events [19]. The use of a single galvo mirror to control all parameters makes it possible to reach sub-millisecond pattern switching speeds that conventional moving gratings and most commercial off-the-shelf SLMs cannot match. Only advanced display technologies like DMDs and ferroelectric SLMs can compete in this regard, but are chromatic (requiring reconfiguration for each different wavelength), highly inefficient due to substantial power loss to unwanted diffraction orders, and size-limited due to the finite display resolution. In contrast to these techniques, SWIFT is achromatic and power-efficient. While we have not demonstrated the use of a second laser due to funding limitations, there are no wavelength-dependent components in the system and no configuration changes are required to change wavelength, which is important when performing high-speed multicolour imaging. SWIFT exhibits no higher order diffraction, no tight focusing and all the optical surfaces insensitive to laser damage. This also makes it more desirable for high laser power applications than fibre-based solutions[20]. Finally, SWIFT’s ability to control polarization is another advantage over DMDs and SLMs, which must either use an electro-optic modulator to rotate the plane of polarization of all beams, or use a so-called “pizza polarizer” consisting of individual waveplates placed conjugate to the entrance pupil of the objective lens. This latter solution requires a particular fixed polarization for each beam, which is problematic in 3D SIM as the center beam must change polarization to match the outermost azimuthally-polarized beams.

In summary, we present SWIFT as a new concept to generate arbitrary interference patterns from a few point sources, with the selection of sources, their intensity, polarization and phase relationships all controlled by a single galvo mirror. In implementing this concept, we have designed a pattern generation device and demonstrated its utility for both 2D and 3D SIM.

## 2 Methods

### 2.1 SWIFT instrument

A 500mW 473nm DPSS laser (LaserQuantum Gem 473) was projected through a variable beam expander before entering a three-way beamsplitter array composed of half wave plates (Thorlabs FBR-AH1) and polarizing cube beamsplitters (Thorlabs PBS051 mounted on FBTC mounts). The reflected s-polarized beams are combined into a set of three closely-spaced parallel rays using pick-off mirrors (Thorlabs FBT-P01) before passing through a small Glan Laser prism (Thorlabs GTH5M-A) to improve the polarization purity. An illustration of this beamsplitter array can be seen in Fig. 2.

The three beams are reflected from a pick-off mirror (Thorlabs BBD1-E02) into a 4*×* demagnifier consisting of a 180 mm focal length achromat (Thorlabs AC508-180-A-ML) and microscope objective (Olympus PLN4X) to strike a galvo mirror (Scanlab Dynaxis 3S). The demagnification ratio must be chosen to minimise the beam diameters on the galvo mirror, within the limits set by lens working distances and the need to fit the resulting foci on the mirror array. Because the beam diameters on the galvo mirror are on the order of one hundred microns, when projected through the ∼200 mm focal length lens before the mirror array, paradoxically the size of the resulting ‘focus’ is significantly larger than that of the ‘collimated beam’ reflected from the galvo mirror. The focus diameter *x* can be estimated from the Rayleigh criterion *x* ≈0.61*λf/D*, where *λ* is the wavelength, *f* is the focal length of the lens and *D* is the aforementioned beam diameter. In practice, 4*×* demagnification is acceptable.

The beams are focussed onto a mirror array by an achromatic lens (Thorlabs ACT508-200-A-ML placed in a telecentric scanning configuration). This mirror array consists of nine custom-made kinematic mounts with three fine adjusters (Thorlabs F2ES8) per mount, using super-elastic nitinol wire as the source of spring tension. Each platform is tapped with an M2 hole in the middle, within which a matching set screw is located; a mirror (Edmund Optics 87-366) is glued on the screw head using thread locking glue. The axial position of the mirrors can therefore be adjusted using the set screw, which is important when using a diode laser with coherence length as short as a few tens of microns. For our prototype, we use a Diode-Pumped Solid State (DPSS) laser with a more practical ∼2 mm coherence length. Thus, the mirrors are coarsely aligned to have a 1 mm offset along the optical axis between each row to compensate for the path-length differences between the three laser beams introduced by the half wave plates, i.e. the top and middle beams travel through 2 and 1 more half wave plates than the bottom beam respectively. An exploded diagram of this mirror array can be seen in Supplementary Fig. 4.

Optionally a quarter wave plate cut from a polymer sheet (Meadowlark BQ-200×200-0473) can be placed in the correct orientation in front of each mirror to control the linear polarization. One of the challenges for SIM is that at high numerical aperture, efficient interference between two beams requires both to have s-polarization due to the vectorial nature of light. In our case, the laser beam was introduced with a horizontal polarization. For the 6 beams reflecting off the side columns of the mirror array, the majority of the power (75 %) is s-polarized and therefore a high contrast interference contrast can be obtained. However, the beams reflecting off the middle column of the mirror array have p-polarization. To rotate polarization, an approx. 5 *×* 20 mm^2^ stripe was cut from the quarter-wave plate sheet with a 45° angle from the fast axis, and placed in a custom-made holder in front of the middle column of the mirror array. As the laser beams pass through the quarter-wave plate twice before and after reflecting off the mirrors, the polarization was changed from horizontal to circular and then to vertical, i.e. s-polarization when the beams reach the microscope.

The kinematic mounts are adjusted to steer the beam from each mirror so that it passes through the center of the entrance pupil of a desired lens in a miniature lens array. This lens array consists of 7 achromatic lenses (Edmund Optics 63-714, 4 mm diameter, 6 mm focal length) in a hexagonally close-packed array. These bring the light from each beam to a tight focus at the back aperture of a well-corrected large-aperture lens (Thorlabs TTL200MP), forming an interference pattern at its image plane. The interference fringes are projected onto the image plane of a microscope, which relays the pattern onto the sample.

### 2.2 Alignment procedure

To align the SWIFT modules, light is first aligned through the beamsplitters. Two mirrors are used to steer the laser to the axis defined by the beamsplitter centers; the half-wave plates are rotated to provide roughly equal light intensity from each beamsplitter. The beamsplitters are rotated and tilted to coarsely align each beam onto its respective pick-off mirror. The middle beam is used as a reference to align the rest of the system; it strikes the D-mirror which is adjusted so that the beam hits the center mirror of the kinematic mirror array when the galvo is in a neutral position. The tilt and rotation of the remaining beams is adjusted using the individual pick-off mirrors so that the beams strike the mirrors above and below in the center column.

Once the beams are centered on the correct mirrors in the array, the kinematic adjusters behind the mirrors are used to back-project the light through the optical system to the desired miniature lens, and finally to the microscope. Custom made beam blockers are placed in front of the mirror array to only allow one beam to pass for individual adjustment of the mirrors. A layer of fluorescent nanoparticles is used to observe the illumination pattern. The kinematic adjusters are fine tuned so that the centre of the field of view is evenly illuminated. The galvo is then rotated to a positive (or negative) angle so that the beams hit the the mirrors on the left (or right) column. The kinematic adjustment are repeated for the side mirrors. After the above procedure, the system is ready for 2-beam SIM with the middle beam blocked by the custom made beam blocker.

In order to generate accurate 3-beam interference for 3D-SIM, the 2 side laser beams must follow a symmetrical path about the centre beam. This means that all 3 beams need to exit the beam splitter in parallel and the centre beam must pass the centre mini-lens. Any asymmetry in the beam direction can cause a “beating” effect in the resulting interference pattern, significantly degrading the reconstructed images. Therefore, the steer of the laser beams needs to be fine tuned based on the generated interference pattern. This is an iterative process as the minuture mirrors need to be adjusted accordingly. The quality of the interference pattern can be directly observed from the image of the nanoparticles. A good 3-beam interference pattern should have even fringe contrast across the field of view while an asymmetric interference pattern would exhibit alternating bands of fringes and diffraction limited nodes.

Finally, the galvo is calibrated to obtain the voltage setting for demanded phase shifts. To do so, the galvo is scanned in fine steps near the 3 angles corresponding to the 3 mirror columns. The correspond phase shifts are estimated using the home-made SIM reconstruction algorithm from the captured images of fluroescent nanoparticles.

### 2.3 SIM setup

To incorporate SWIFT into a SIM microscope, the interference fringes are placed at the image plane and relayed to the sample via the tube lens (Thorlabs TTL200MP), dichroic mirror (Semrock Di03-R473-T1-25×36, specified *<* 1*λ* peak-to-valley wavefront error) and microscope objective (100*×* 1.5 NA TIRF Olympus UPLAPO100XOHR). All components are mounted in an Olympus IX73 inverted microscope frame, with a custom-made deck insert. For 3D SIM measurements, the focus position was controlled automatically using a stepper motor secured to the fine focusing knob of the microscope (Prior Scientific PS3H122R with Proscan III controller).

To ensure the image of the foci lies just within the back aperture of the objective, a Galilean beam expander combining two lenses (Newport KPC067 and Thorlabs LA1727-AB) is applied between the two TTL200MP lenses to adjust the effective period of the fringes. Images of the sample are projected back through the objective and dichroic, through an emission filter (Semrock BLP01-473R-25) and projected onto a high-speed high-performance camera (Photometrics Kinetix) with the in-built tube lens of the microscope frame.

### 2.4 Sample preparation

To prepare the layer of fluorecent nanoparticles, the suspension of nanoparticles (Fluoresbrite YG Carboxylate microspheres 0.1 μm) was diluted, dropped on to a coverslip and spreaded using the pipette tip before let dry. Afterwards, a few drops of optical adhesive (Norland 81) were dropped onto the beads and then covered by a microscope slide with another 2 slides as spacers between the coverslip and the covering slide, resulting in a 1 mm layer of adhesive on the beads after being cured using ultraviolet exposure. The beads were then imaged through the coverslip.

U2OS FlipIn Trex cells were cultured in DMEM (Corning) supplemented with 10% fetal bovine serum (Gibco) and 1% penicillin–streptomycin (Gibco) at 37 ^*°*^C with 5% CO_2_. Cells were trypsinised and plated on #1.5 coverslips coated with fibronectin (Sigma, F1141, 50 μg / ml in PBS), for 2 h at 37 ^*°*^C in DMEM-10% serum. Cells were screened for mycoplasma using MycoAlert Mycoplasma Detection Kit (Lonza). The medium was removed and the cells were washed twice with warm PBS and fixed with 4% paraformaldyhde. After a further two PBS washes, 100 μl of 1 μl in 1 ml phalloidin-iFluor 488 (Abcam) with 1% BSA was added and left for 90 minutes. Cells were then washed again twice with PBS and mounted onto slides using Fluoromount-G (Invitrogen).

