## Supplementary information for "Synthesis of arbitrary interference patterns using a single galvanometric mirror, and its application to Structured Illumination Microscopy"

#### 1 Adjusting focal separation

The miniature lens array determines the angle of the interfering plane waves, but in some cases it is desirable to tune these angles by changing the magnification of the foci created by the lens array. In SIM for example, the image of these foci should lie just within the back aperture of the microscope objective lens. To tune the spacing of the foci, a lens system can be placed between the two lenses that relay the foci to the objective back aperture. Three example configurations (for spacings of 4.6 mm, 5 mm and 7 mm) are given in supplementary table 1-3, and their layouts can be seen in Supplementary Fig. 1.

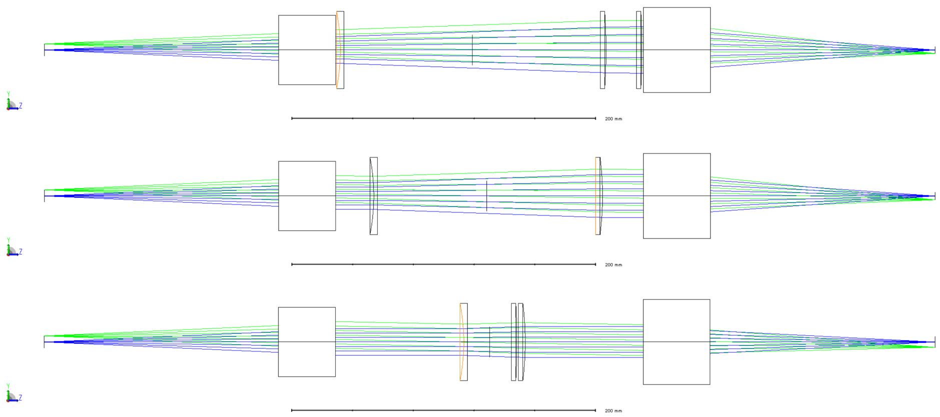

**Supplementary Figure 1** Layouts for adjusting focal separation. Top: 4.6mm focus separation. Middle: 5mm focus separation. Bottom: 7mm focus separation. Total track length is 432.3mm in all cases.

| *Lens* | *Distance to next lens surface* |
|---|---|
| Miniature lens focus | 154.1mm |
| ThorLabs TTL200MP | 3.0mm |
| Newport KPC067 | 168.6mm |
| ThorLabs LA1727-A | 20.0mm |
| ThorLabs LA1779-A | 1.0mm |
| ThorLabs TTL200MP | 148.0mm |
| Objective back aperture | |

**Table 1** 4.6mm focal separation

| *Lens* | *Distance to next lens surface* |
|---|---|
| Miniature lens focus | 154.1mm |
| ThorLabs TTL200MP | 25.0mm |
| Newport KPC067 | 143.5mm |
| ThorLabs LA1725-A | 26.9mm |
| ThorLabs TTL200MP | 148.0mm |
| Objective back aperture | |

**Table 2** 5mm focal separation

| *Lens* | *Distance to next lens surface* |
|---|---|
| Miniature lens focus | 154.1mm |
| ThorLabs TTL200MP | 83.1mm |
| Newport KPC067 | 28.8mm |
| ThorLabs LA1779-A | 1.0mm |
| ThorLabs LA1725-A | 76.4mm |
| ThorLabs TTL200MP | 148.0mm |
| Objective back aperture | |

**Table 3** 7mm focal separation

#### 2 Custom parts

Several custom parts are needed to assemble SWIFT, primarily because off-the-shelf optomechanics were too bulky. CAD models are all available either upon request or from https://www.imperial.ac.uk/rowlands-lab/.

##### 2.1 Beamsplitter fiberbench

The purpose of the fiberbench system is to split the incident beam into three beams of identical polarization but with an arbitrary split of intensity. The system used ThorLabs fiberbench components for reasons of cost and compactness, but required a custom-machined baseplate as suitable commercial alternatives were unavailable.

##### 2.2 Miniature lens and pick-off mirror assembly

The seven miniature lenses are sandwiched between two custom-machined aluminium plates secured inside a lens tube. Before mounting, the lenses were first bundled together using a rubber band to achieve an accurate hexagonal packing. This step should not be neglected, since any inaccuracy in the mounting process can result in asymmetry in the hexagonal arrangement and consequently a beating effect in the 3-beam interference pattern.

##### 2.3 Kinematic mirror array

Because of the limited field of view of the lens between the galvo mirror and the miniature mirrors, the mirror array was custom-built in order to fit a $3\times3$ mirror array within a 25 mm field of view. Each mirror was mounted on a custom-designed platform that could achieve tip, tilt and piston actuation via a series of set screws; the piston was used for path-length matching in order to optimize the interference contrast and tip-tilt allows each beam to address a particular miniature lens.

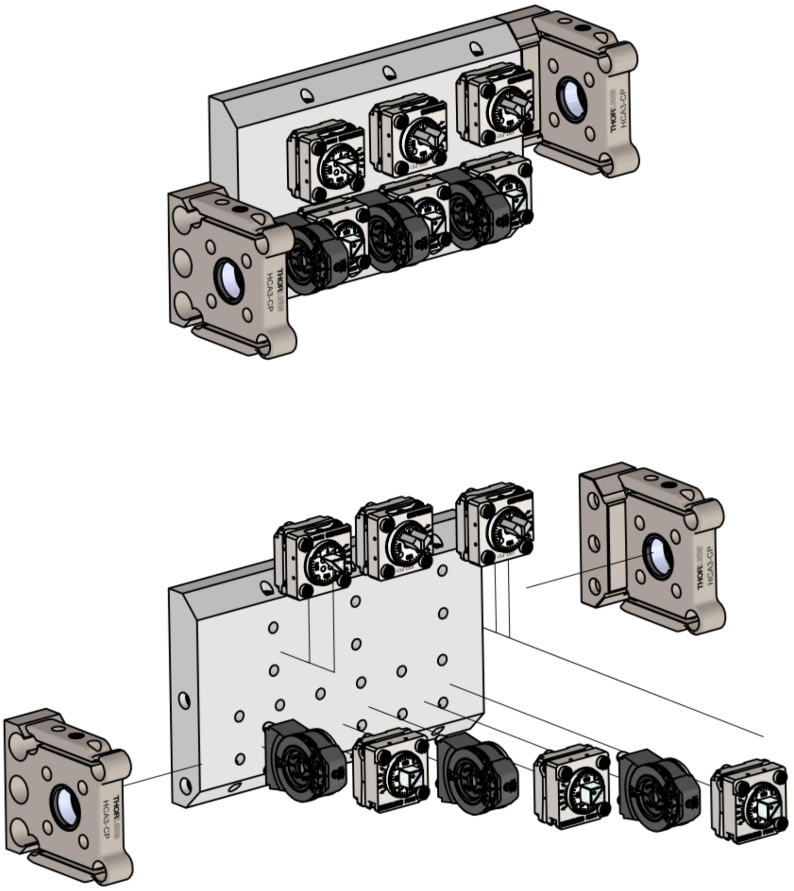

**Supplementary Figure 2** Beamsplitter construction. Top: Laser enters through the bottom cage mount and passes into three sequential pairs of half wave plates and polarizing beamsplitters. These can be tuned to pick off desired fractions of the primary laser beam at each beamsplitter. The pick-off mirror above each beamsplitter allows the beams to be oriented parallel to each other but with a 2 mm offset between them. Bottom: Exploded diagram showing each component and its assembly.

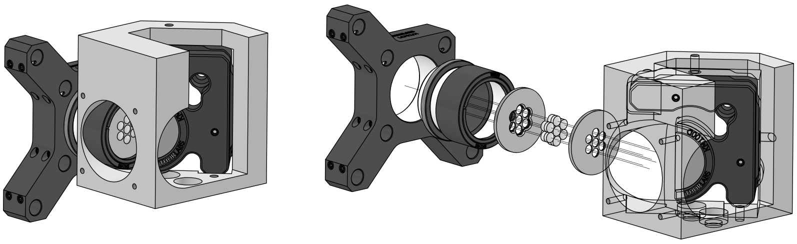

**Supplementary Figure 3** Miniature lens array and pick-off mirror. Left: Light enters from the left; there are standard #4-40 taps for connection to a 30 mm cage system. The light is reflected from a pick-off mirror to the rest of the optical system through the cutout to the right. Returning light illuminates one of seven microlenses, bringing it to a focus. Right: Exploded diagram showing the individual components.

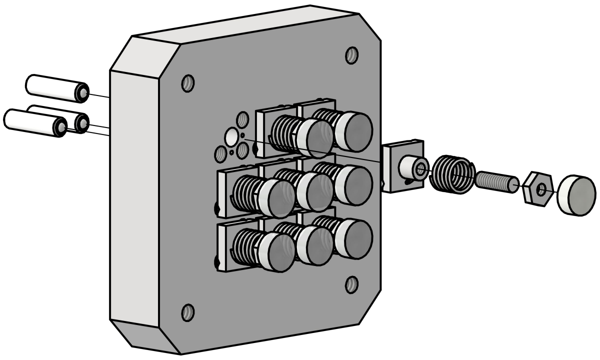

**Supplementary Figure 4** CAD model of the mirror array, with one mirror and its mount shown in exploded layout. Each mirror is mounted on a kinematic platform, which permits adjustment of tip, tilt and axial position. Each custom-machined kinematic platform is secured to the base with superelastic nitinol wire as a spring, with three 0.2 mm pitch set screws to adjust tip, tilt and tension. The mirror itself is mounted on a platform attached to a spring-tensioned set screw which can translate to match the path lengths of each laser beam. The entire assembly is compatible with standard 30 mm cage systems.

### 3 SIM reconstruction software

The SIM images were reconstructed in the following steps using a home-made MATLAB program partly based on open source algorithms [1, 2].

- **Preprocessing**
  - When performing fast 2D-SIM imaging, there can be microscale movement of the sample which degrades the reconstructed image. To minimise the influence of such movement, the raw frames were first compared to the first frame to detect and cancel the shifts of the sample image using a sub-pixel image registration algorithm [3].
  - For 3D-SIM imaging, the raw frames were rearranged to form 15 3D-images with 3 orientation and 5 phases.
  - The 2D or 3D images were normalized to compensate for the different illumination powers of the three orientations. To do so, all images with the same illumination pattern orientation were divided by the same factor of their average counts.
- **Parameter estimation** The illumination spatial frequencies, the phase shifts and the amplitude of each frequency component were determined *a posteriori* from the pre-processed images independently for the 3 orientations as following.
  - An average of all images were calculated as an estimate of the sample image with homogeneous illumination, i.e. the central frequency component. This is a good estimate for the 2D images. The average 3D image would have an extra modulation in the axial direction, which however does not significantly affect the parameter estimation as the spatial modulations to be estimated are lateral.
  - The illumination spatial frequencies were determined by cross-correlating of Fourier tranforms of the pre-processed images and the average image. In the 3D case, the first order spatial frequencies were estimated and the second order were calculated by doubling the first order frequencies.
  - To estimate the phase shifts and amplitudes of the frequency shifted image components of the 2D images, each pre-processed image was considered as a linear combination of 3 components: the average image (i.e. the central frequency component), the product of the average image and a 2D cosine pattern with the illumination spatial frequency (i.e. a frequency shifted image component with no phase shift), and the product of the average image and corresponding sine pattern (i.e. a frequency shifted image component with $\pi/2$ phase shift). The ratio of the components were determined using linear regression (MATLAB mldivide). The phase shifts of the illumination pattern and the amplitude of each frequency shifted image component were calculated from the ratio of the components based on the angle addition theorem. Note that the amplitudes obtained here were reduced by the OTF.
  - Similar calculations were applied to 3D images, except they have five components with two spatial frequencies.
- **Reconstruction**

- A theoretical OTF was estimated assuming NA = 1.48. The 3D OTF was estimated using the algorithm in [4].
- The different frequency-shifted image components and the central frequency component were separated based on the estimated parameters.
- All components from three orientations were recombined after applying a generalized Wiener filter based on the calculated OTF and corresponding spatial frequency shifts.
- A notch filter [1] was applied to the 3D images to reduce artifacts.
- Images were thresholded to further reduce artifacts.

[4] Manton, J. Debye diffraction code. https://github.com/jdmanton/debye_diffraction_code.
